# New single-molecule imaging of the eisosome BAR domain protein Pil1p reveals filament-like dynamics

**DOI:** 10.1101/092536

**Authors:** Michael M. Lacy, David Baddeley, Julien Berro

## Abstract

Molecular assemblies can have highly heterogeneous dynamics within the cell, but the limitations of conventional fluorescence microscopy can mask nanometer-scale features. We have developed a novel, broadly applicable, fluorescent labeling and imaging protocol, called Single-molecule Recovery After Photobleaching (SRAP), which allowed us to reveal the heterogeneous dynamics of the eisosome, a multi-protein structure on the cytoplasmic face of the plasma membrane in fungi. By fluorescently labeling only a small fraction of cellular Pil1p, the core eisosome BAR domain protein in fission yeast, we visualized whole eisosomes and, after photobleaching, recorded the binding of individual Pil1p molecules with ~20 nm precision. Further analysis of these dynamic structures and comparison to computer simulations allowed us to show that Pil1p exchange is spatially heterogeneous, supporting a new model of the eisosome as a dynamic filament.

**Abbreviations used:**

BAR: Bin/Amphiphysin/Rvs domain
FRAP: Fluorescence Recovery After Photobleaching
mEGFP: monomeric Enhanced Green Fluorescent Protein
SiR647: silicon-rhodamine 647
TIRF: Total Internal Reflection Fluorescence

## Introduction

Fluorescence microscopy including methods such as FRAP (Fluorescence Recovery After Photobleaching) have been invaluable for characterizing cellular organization and dynamics at the micrometer scale. However, it has been particularly challenging to characterize spatial heterogeneities inside diffraction-limited zones and dynamics within individual multi-molecular assemblies in live cells.

The eisosome is a multi-molecular assembly on the cytoplasmic face of the plasma membranes of fungi, consisting of a stable scaffold of proteins clustered on a small invagination of membrane (Douglas and Konopka 2014; (Strádalová et al. 2009;(Walther et al. 2006;(Malínská et al. 2003), whose various functions in cell membrane regulation remain unclear (Kabeche, Howard, and Moseley 2015;(Aguilar et al. 2010;(Fröhlich et al. 2014;(Kabeche et al. 2015). Fission yeast eisosomes are linear (50 nm wide and 1-2 μm long), while budding yeast eisosomes appear as diffraction-limited puncta. The main protein component of the eisosome in fission yeast, Pil1p, contains a Bin/Amphiphysin/Rvs (BAR) domain which facilitates its organization *in vivo* (Ziółkowska et al. 2011;(Olivera-Couto et al. 2011) and its oligomerization *in vitro* (Karotki et al. 2011;(Kabeche et al. 2011), features conserved in budding yeast Pil1 and other BAR-domain containing proteins.

Eisosomes are essentially immobile and exhibit no dynamics in Fluorescence Recovery After Photobleaching (FRAP) experiments on timescales of 10 to 20 minutes (Kabeche et al. 2011;(Walther et al. 2006), which has led to the conclusion that the eisosome is a static microdomain. However, recent reports suggest that eisosomes may be more dynamic than previously believed: in cells lacking a cell wall, eisosomes disassemble in response to increased turgor pressure (Kabeche, Howard, and Moseley 2015), and dynamic sub-populations of Pil1 oligomers exist on the membrane near eisosomes (Olivera-Couto et al. 2015).

In this report, we show that the eisosome ends are dynamic while its core is stable, as in an oligomeric filament. To demonstrate this result, we developed a novel fluorescence imaging strategy to monitor single-molecule dynamics in live cells, called Single-molecule Recovery After Photobleaching (SRAP). By labeling only a small fraction of Pil1p molecules, we visualized whole eisosomes in the first few frames of a movie, and after photobleaching we observed isolated Pil1p molecules re-appearing specifically at the ends of eisosomes. By comparison with computer simulations, we demonstrate that our data support a model of the eisosome as a dynamic filament.

## Results

### Single-molecule recovery after photobleaching of Pil1p

To perform our single molecule experiments in live cells, we sparsely labeled Pil1p by fusing a SNAP-tag to the protein C-terminus and incubating cells with relatively low concentration (0.5 μM) of benzylguanine-conjugated silicon-rhodamine dye (SiR647) (Keppler et al. 2002;(Lukinavičius et al. 2013). This protocol yielded sufficient density of Pil1p-SNAP labeled with SiR647 (referred to as Pil1p-SiR) to visualize long linear eisosomes on the cell membrane (Supplemental Figure S1A), similar to structures observed in cells expressing Pil1p-mEGFP (Supplemental Figure S1B-C), and very low non-specific labeling (Supplemental Figure S2).

After about 5 seconds of imaging under low-power TIRF illumination (~20 W/cm^2^), the fluorescently-labeled eisosomes visible in the first few frames photobleached. Because TIRF imaging only illuminates molecules within ~500 nm above the coverslip, unbleached Pil1p-SiR molecules in the cytoplasm beyond the TIRF field may diffuse into the illumination field in later frames of the movie (Figure 1 A and B, and see Supplemental Movie 1). Since only a small fraction of Pil1p molecules were fluorescently labeled, fluorescence re-appeared as isolated diffraction-limited spots corresponding to single Pil1p-SiR molecules. Intensity traces of sites of recovery revealed stepwise decreases (corresponding to photobleaching or unbinding) and increases (corresponding to binding of single Pil1p-SiR molecules) over the time course of the movie (Figure 1C), characteristic of single fluorescent molecules. In addition, recovery spots were immobile, suggesting that they were not freely diffusing on the membrane surface and indeed corresponded to fluorescent Pil1p-SiR incorporated into eisosomes. This sparse labeling and imaging strategy, which we call Single-molecule Recovery After Photobleaching (SRAP), revealed that new Pil1p molecules bind at eisosomes within a few seconds after initial photobleaching of the labeled structure.

**Figure 1.**
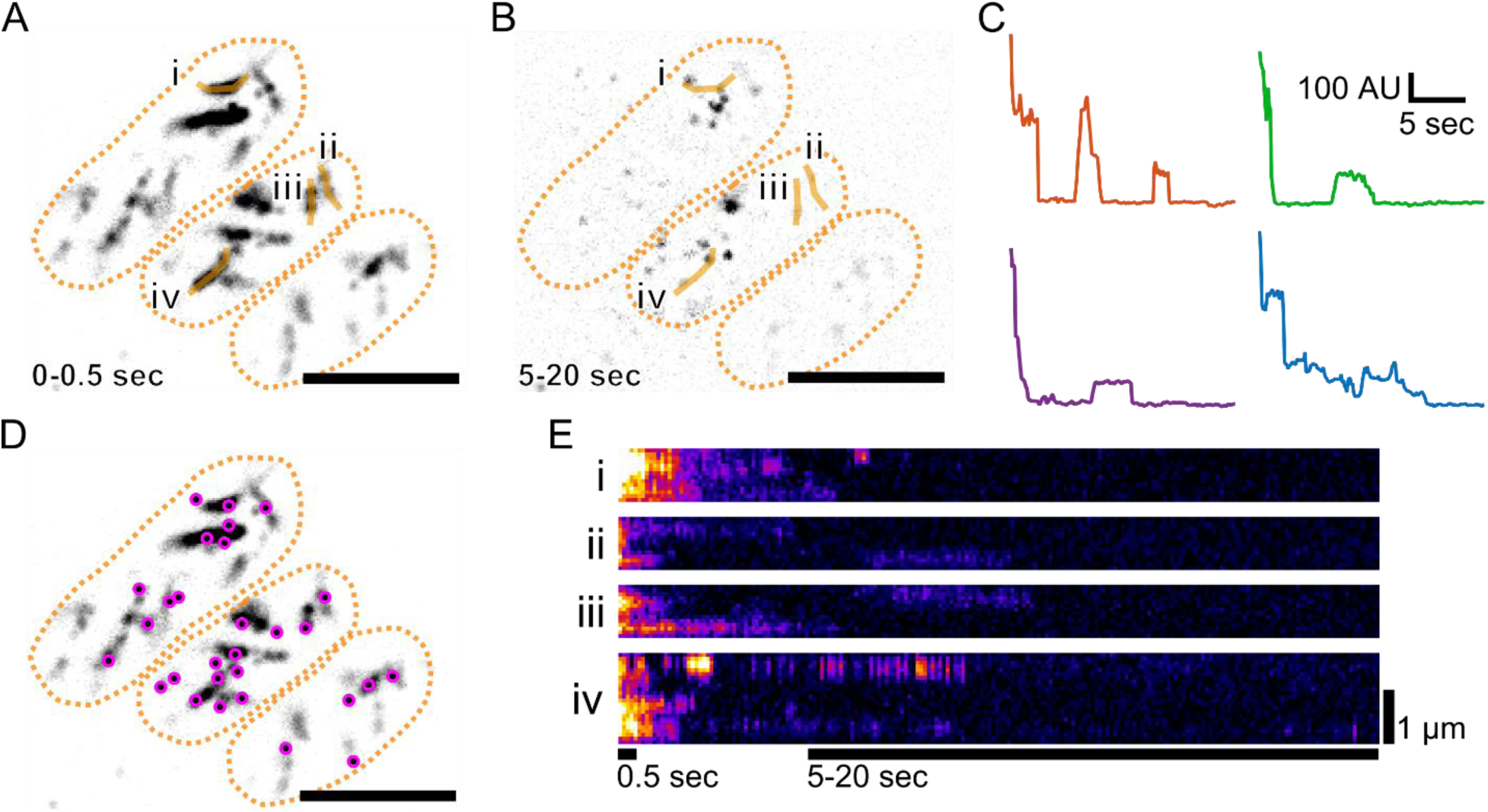
Single-molecule recovery after photobleaching of labeled eisosomes. Cells expressing Pil1p-SNAP were labeled with SNAP-SiR647 at 0.5 μM for 15 hours, washed and imaged in TIRF. (A) Average intensity projection of the first 5 frames (0.5 seconds) of a movie reveals linear eisosomes. (B) Maximum intensity projection of frames 50-200 (5 to 20 sec) of the same movie shows single molecule recovery events. Cell outlines are drawn in orange dashed lines. Orange lines are the line traces used for the kymographs in E. (C) Example intensity traces of recovery spots show stepwise photobleaching and single-molecule recovery of Pil1p-SiR. The intensity of a 3-pixel diameter circle was measured across all frames of the movie after subtracting the median-filter background. Intensity traces were processed with a Chung-Kennedy filter to highlight discrete intensity steps. (D) The positions of single-molecule recovery events (magenta spots, maxima detected in image B) are mapped on the eisosomes (same image as A). (E) Kymographs of line traces along eisosomes as labeled in A and B, with bars indicating the time spans for the projection images. A, B, D scale bar: 5 μm.

We measured the lifetimes of these fluorescence recovery events and fit the distribution of lifetimes with an exponential curve. The disappearance rate, 2.6 ± 0.2 sec^−1^, is faster than the overall rate of photobleaching in the images, 0.48 ± 0.03 sec^−1^ (95% confidence intervals, see Supplemental Figure S3), suggesting that Pil1p-SiR molecules are not only photobleaching but also unbinding from the eisosomes. By subtracting the photobleaching rate from the spots' disappearance rate, we estimate the observed unbinding rate of Pil1p to be approximately 2.1 ± 0.2 sec^−1^, consistent with the findings of Olivera-Couto *et al.* (2015). We do not attribute these recovery events to fluorophore blinking, as SiR647 displays less blinking behavior compared with other derivative dyes (Uno et al. 2014) and usually requires high laser intensity or additives to enhance dark state switching. Therefore, we conclude that Pil1p is indeed undergoing fast single-molecule exchange at eisosomes, binding and unbinding even in the absence of large-scale eisosome remodeling.

### Pil1p recruitment is not uniformly distributed

Further inspection of the recovery events suggested that eisosome ends are hotspots of Pil1p exchange (Figure 1D). Kymographs of lines drawn along eisosomes showed that fluorescence signal at eisosome ends persisted longer and recovered after photobleaching more frequently than along the interior (Figure 1E). To precisely calculate the distance of SRAP spots to the eisosome end, we determined the position of each spot with super-resolution localization and determined the position of each eisosome end by fitting a sigmoidal curve to the intensity profile traced along the eisosome end in initial frames (see Materials and Methods and Figure 2A). We found 92% of SRAP spots were within 250 nm from their corresponding eisosome end, with an average position of 97 ± 119 nm (mean ± S.D., N = 191 spots in 20 cells, Figure 2B).

**Figure 2.**
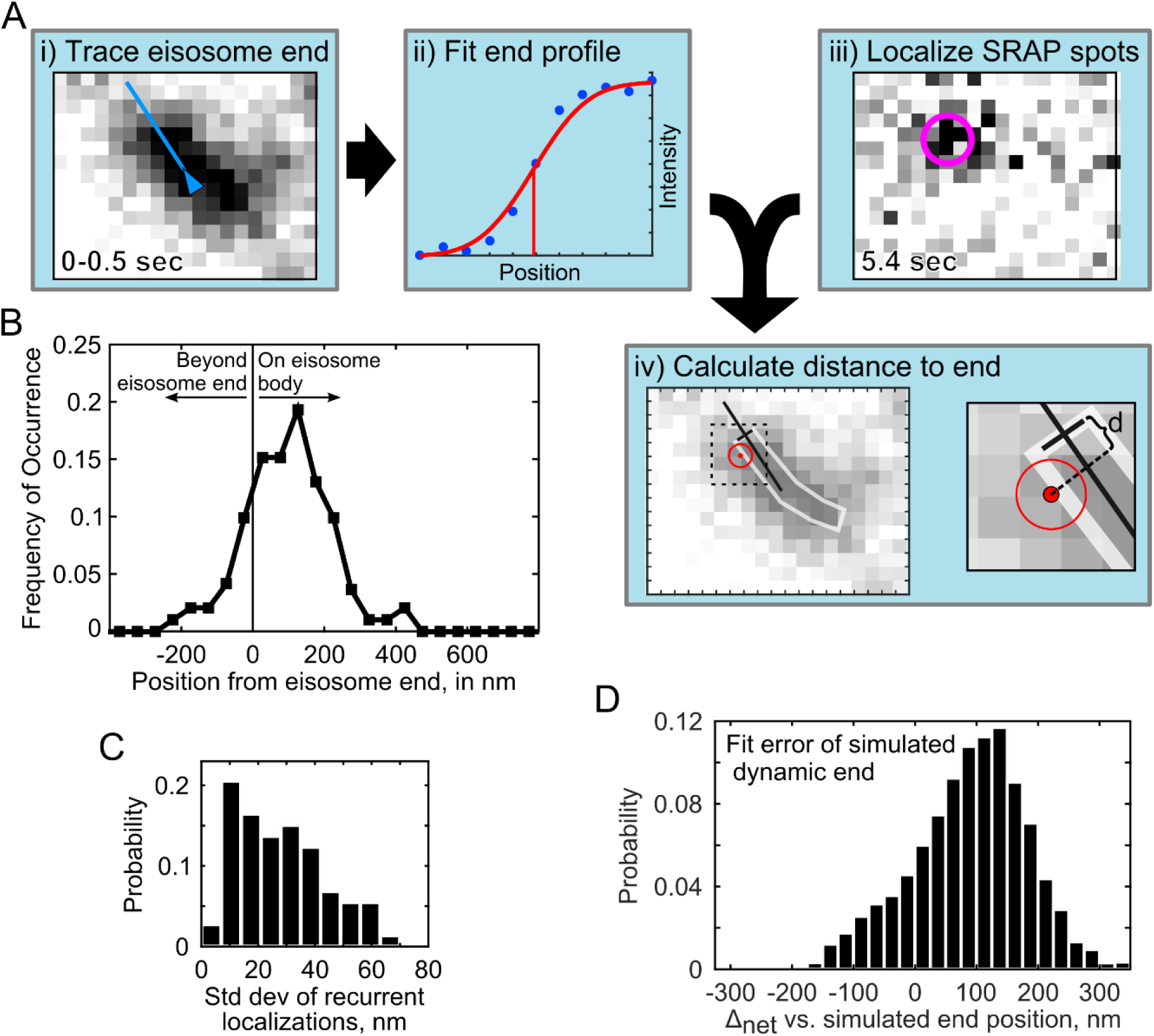
Image analysis for localization of SRAP spots at eisosomes. (A) Schematic of the measurement of distance to eisosome end: (i) The end of an eisosome is traced in the average projection of the first 5 frames (0-0.5 sec); (ii) the line intensity profile of the eisosome end is fitted to determine the position of the diffraction-limited end (red line); (iii) a SRAP spot position is determined with the PeakFit plugin for ImageJ in the movie frame when it appeared, and super-resolution localizations from multiple frames are averaged to calculate the position of the SRAP event; (iv) the distance *d* is calculated from the SRAP spot along the eisosome line trace to the end; in (i, iii, iv), one image pixel is 70 nm. (B) measured SRAP spot positions relative to the eisosome end, average 97 ± 119 nm S.D. (N = 191 spot/filament pairs across 20 cells). (C) Standard deviation calculated for each SRAP spot that included multiple localizations, average 27.9 ± 15.9 nm S.D. (N = 73 sets). (D) Simulated errors of the fitting of sparsely labeled, dynamic eisosome ends. Mock eisosome end intensity profiles (as in A.ii) were generated according to a 1% labeling efficiency, with a single extra emitter added to the end position and fitted as described in Materials and Methods. Average difference between the fitted end position and the simulated true end is 89.3 ± 94 nm S.D. (N = 5000 simulations).

To more clearly interpret this distribution of SRAP positions, we simulated datasets based on hypothetical models for Pil1p recovery dynamics. In a first model (referred to as the “uniform model”) we assumed that binding events occur uniformly along the eisosome (Figure 3A, blue). The simulations included noise terms to mimic the uncertainty in the localizations for the SRAP spots and the eisosome end (Figure 2C and Supplemental Figure S4C), and took into account the distribution of eisosome lengths measured experimentally (Supplemental Figure S1C). The distribution of the measured data was in poor agreement with the uniform model (Figure 3A-B).

**Figure 3.**
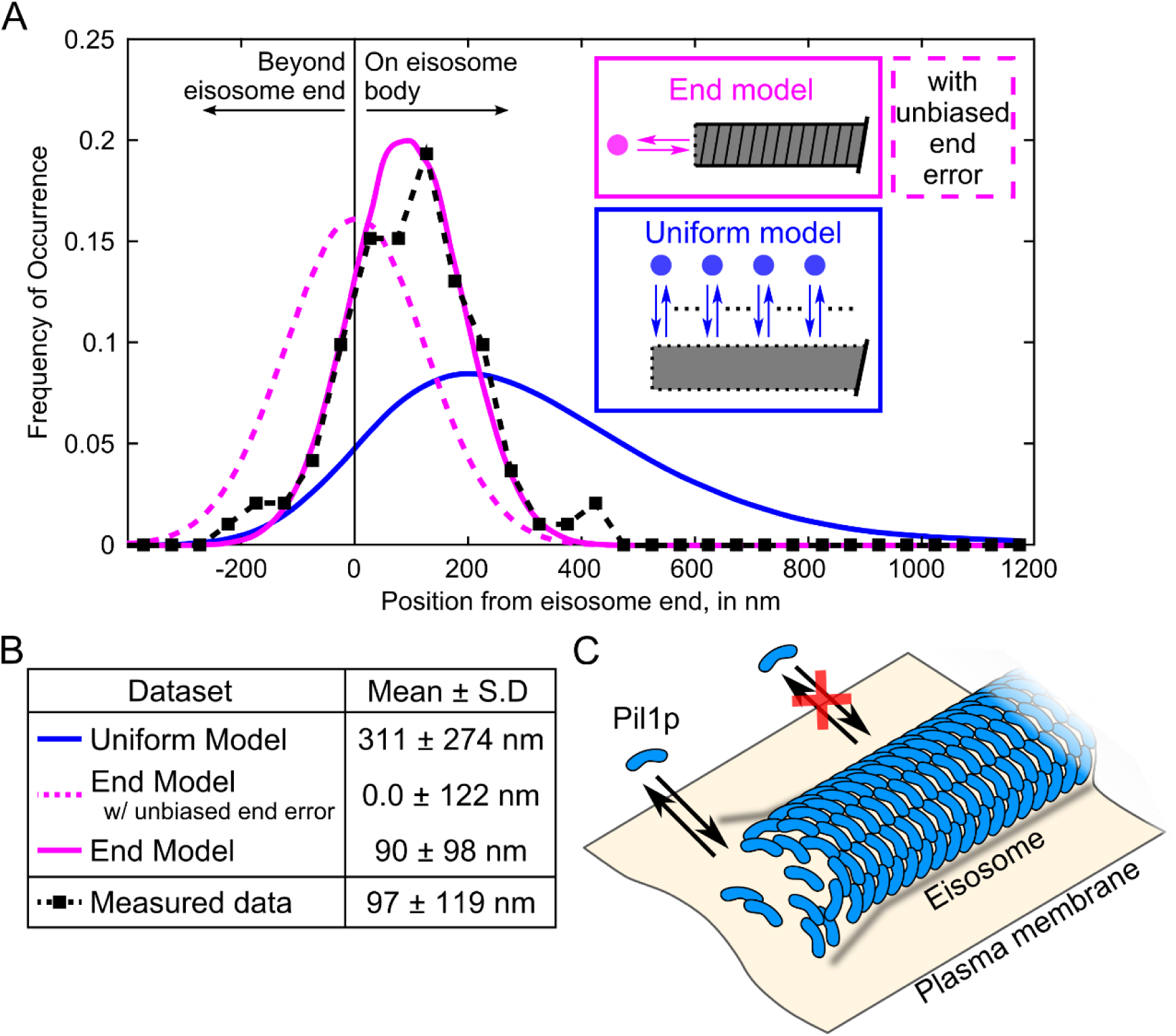
Single-molecule recovery of Pil1p-SiR occurs at eisosome filament ends. (A) Probability distributions of measured distances (black squares, N = 191 spot/filament pairs across 20 cells) and simulation results (uniform dynamics model, blue; end dynamics model, magenta, N = 275,000 runs for each tested model). The simulation models are illustrated in schematic form, the “uniform model” where recovery events are distributed uniformly along the eisosome length (blue), or the “end model” where recovery events are confined to eisosome ends (magenta). For the end model simulations, the dashed line represents simulations using an unbiased Gaussian noise distribution for eisosome end localizations (as in Supplemental Figure S4E), and the solid line represents simulations using the noise predicted from a dynamic end (as in Figure 2D). (B) Table of mean and standard deviation of distributions for simulated datasets and measured SRAP spots. The difference between the measured data and the end model with dynamic end noise is not statistically significant, by two-sample Kolmogorov-Smirnov test (p = 0.39). (C) Model of the eisosome as a dynamic filament: Pil1p subunits assembled into a filament on the cytoplasmic surface of the plasma membrane are free to associate and dissociate at the ends but not at the interior.

In a second model (referred to as the “end model”), we assumed that binding of new Pil1p occurred only at the eisosome ends, as in a dynamic filament. The distribution of simulated recovery positions followed a shape similar to our experimental data but with a mean of 0 ± 122 nm (Figure 3A-B, dashed magenta). The slight bias of the SRAP spot localizations towards the interior of the eisosome (97 ± 119 nm) seems in contradiction with a model of dynamics strictly confined to the end. This shift cannot be explained by an error in our spot localization precision, as the standard deviation of positions calculated for recurrent localizations at a given SRAP site was 27.9 ± 15.9 nm (Figure 2C). We wondered if this shift could be explained by the accuracy of our localization of eisosome ends and if a dynamic end could introduce a systematic bias.

### Localization accuracy of sparsely labeled, dynamic eisosome ends

We evaluated the accuracy of our eisosome end localization method using simulated data mimicking sparsely labeled linear filaments, with densities similar to our experimental data (Supplemental Figure S4A). First, we found that determining the sub-diffraction-limited position of the end of the sparsely labeled eisosome by fitting the intensity profile with an error function – a model that assumes a continuous distribution of emitters – typically over-estimates the true position of the edge of the labeled region towards the exterior of the structure by 50 to 150 nm, depending on the number of fluorophores contributing to the intensity trace. But, because the eisosomes are sparsely labeled, the distance between the true end of the eisosome and the fluorescently labeled Pil1p closest to the end counteracts this bias (Supplemental Figure S4A-B). In sum, in our simulations corresponding to 1% labeling efficiency, the average error of the fitted eisosome end position is only 3.2 ± 119 nm (S.D., Supplemental Figure S4C).

However, because we used intensity profiles extracted from a projection image averaged over a short time, if the recruitment of Pil1p is indeed localized to the eisosome end, then any new labeled molecules that bind during the recording time would skew the intensity profile towards the end (Supplemental Figure S4D). Indeed, in kymographs of sparsely labeled eisosomes (Figure 1E), the signal at eisosome ends persisted longer than the signal along the filament body. We simulated this effect by adding a single extra emitter at the true end position before fitting the intensity profile, resulting in a net error of 89.3 ± 94 nm (Figure 2D, and Supplemental Figure S4E-F), mirroring the offset in our measured SRAP spot positions.

### Eisosome ends are specific sites of single-molecule recovery events

When we repeated simulations of the end model for eisosome recovery incorporating this biased localization error for the eisosome end, the result clearly reproduced our experimental data (90 ± 98 nm, Figure 3A-B, solid magenta, *p* = 0.39 by Kolmogorov-Smirnov test). Importantly, the only assumption of this model is that the eisosome end is the specific site of Pil1p binding: the bias in the eisosome end localization arises from the sparse labeling of the sample.

As a putative alternative hypothesis, we considered a model in which Pil1p binding is restricted to a region near the eisosome end rather than strictly confined to the end. Simulations of this “hybrid model” using a 200-nm region at the eisosome end produced a similar distribution of positions (100 ± 103 nm, Supplemental Figure S5, *p* = 0.32 by Kolmogorov-Smirnov test). However, the standard deviation of recurrent localizations at the same SRAP site (27.9 ± 15.9 nm, Figure 2C) indicated that binding events occur at a fixed position on each eisosome, in contradiction with a dynamic 200-nm region. Therefore, we believe the most likely explanation of our experimental data is that Pil1p recruitment occurs at the eisosome ends but the method for determining the end positions of this sparsely labeled dynamic structure introduces a slight systematic bias.

## Discussion

### SRAP reveals heterogeneities at the nanometer scale *in vivo*

The localization of single molecule exchange at the eisosome has previously been unobservable using fluorescent fusion proteins. Our new SRAP method, which relies on partial labeling and photobleaching during imaging, was critical for revealing the behavior of individual protein molecules in the context of the larger eisosome structure. This protocol is the first reported use of SNAP-tag in live fission yeast, and we expect our new SRAP method will be easily applicable to study single-molecule dynamics and heterogeneities in other multi-molecular assemblies in any organism. While similar sparse fluorescence conditions might be achieved by partial photobleaching or photoswitchable proteins, our labeling protocol has the advantage of using organic fluorophores which are brighter and more photostable than fluorescent proteins, providing better single-molecule localization precision.

### Filament model for the eisosome

Our results demonstrate that the eisosome exists in a dynamic steady state with continuous and fast exchange at its ends, even in the absence of perturbation. Models of the eisosome as a static membrane compartment or microdomain (Kabeche et al. 2011;(Karotki et al. 2011;(Walther et al. 2006) would predict monomer exchange to occur uniformly around its edges. Instead we propose a new model for the eisosome as a membrane-bound filament of Pil1p with a stable core and ends in dynamic equilibrium (Figure 3C). Pil1p and other BAR domain-containing proteins have been observed to oligomerize and form filaments and membrane tubules *in vitro,* but it has been unclear to what extent this oligomerization exists *in vivo* or if instead BAR proteins are simply clustered in patches (Adam, Basnet, and Mizuno 2015;(Daum et al. 2016;(McDonald and Gould 2016;(Suetsugu 2016). Recent *in vitro* studies of BAR proteins have proposed that rigid oligomeric scaffolds on membrane tubes can arise without formation of a dense lattice (Simunovic et al. 2016), but the binding dynamics of individual molecules within such scaffolds have not been characterized. Our results are the first observation of any BAR protein scaffold displaying dynamics consistent with a membrane-bound filament *in vivo*.

Our dynamic filament model is consistent with previous FRAP experiments because the few molecules at filament ends that undergo exchange would be virtually impossible to observe with bulk fluorescent imaging since they represent only a very small fraction of the total fluorescent signal. Our model also explains the heterogeneous binding equilibria observed by Olivera-Couto *et al.* (2015). Importantly, a filament model predicts that eisosome remodeling could occur in response to physical or biochemical cues by simply altering the equilibrium of subunits at the eisosome ends by modulating the rates of Pil1p binding or unbinding, just like other cytoskeletal filaments such as actin and microtubules.

## Materials and Methods

### Yeast strains and SNAP labeling

We tagged the pil1 gene at its C-terminus with SNAP-tag (cloned from Addgene Plasmid #29652 pENTR4-SNAPf, inserted into pFA6a vector with KanMX6 selection marker) or mEGFP, in its native locus in a wild-type *S. pombe* strain by homologous recombination (Bähler et al. 1998). Cells were grown at 32° C in liquid YE5S medium to exponential phase (OD_595nm_ between 0.4 and 0.8), then diluted into liquid EMM5S medium and grown for 12 to 24 hours at 25° C before labeling with SNAP fluorophore.

Although the SNAP-tag has been used successfully in a variety of applications (Bosch et al. 2014;(Stagge et al. 2013; (Klein, Proppert, and Sauer 2014), labeling cellular SNAP fusion proteins in live yeast is difficult because the cell wall impedes entry of the fluorophore substrate and because multi-drug exporters prevent its accumulation in the cytoplasm (McMurray and Thorner 2008; (Stagge et al. 2013). These issues may be avoided by enzymatically digesting the cell wall, deleting the multi-drug exporter genes (McMurray and Thorner 2008) or using electroporation to allow a large amount of dye to enter the cells (Stagge et al. 2013). However, such approaches may be problematic if the structure of interest is sensitive to cell integrity, as is the case with the eisosome. To avoid these difficulties, we used a minimally disruptive approach, adding a low concentration of SNAP substrate fluorophore in the media for a long incubation.

To label SNAP-tag protein in live cells, 0.5 mL of cells at OD_595nm_ 0.5 were incubated at 25^o^ C on a rotator in liquid EMM5S media containing 0.1, 0.5, or 2.5 μM of the siliconrhodamine benzylguanine derivative SNAP-SiR647 or SNAP-Alexa647 (SNAP-Cell^®^ 647-SiR, SNAP-Surface^®^ Alexa Fluor^®^ 647, New England Biolabs) for 0.5, 5, or 15 hours. For samples incubated for 15 hours, the cells were initially diluted to OD_595nm_ of 0.1 to avoid over-growing during the incubation time. Cells were washed three times by centrifuging at 900xg for 3 minutes and resuspending in 0.5 mL of EMM5S, then additionally incubated at 25^o^ C for one hour in 0.5 mL of EMM5S, then washed three times again by centrifuging at 900xg for 3 minutes and resuspending in 0.5 mL of EMM5S. Cells were finally resuspended in 50 to 100 μL of 0.22-μm filtered EMM5S to achieve suitable cell density for imaging.

We estimated the extent of labeling by dividing the total intensity of cells in the first frame by the mean pixel intensity of the late-appearing single molecule spots to determine the number of fluorophores per cell. We then determined the fraction of labeled Pil1p-SNAP molecules by dividing the number of fluorophores per cell by the total number of Pil1p molecules expressed in fission yeast cells (Carpy et al. 2014) as reported in PomBase (Wood et al. 2012; McDowall et al. 2015). Although there is significant uncertainty in this estimation, the samples we used for SRAP analysis consistently had labeling efficiencies between 1 and 3%.

Our protocol still requires use of a cell-permeable fluorophore conjugate, as incubation with SNAP-Alexa647 yielded poor labeling (Supplemental Figure S2, B and C). Incubation with 2.5 μM of SiR647 for 15 hours achieved a higher density of labeled Pil1p-SiR (10% or more, Supplemental Figure S1A), but short incubations yielded only sparse labeling with greater cell-to-cell variability (Supplemental Figure S2A).

### Microscopy

Live cells were imaged on 25% gelatin pads in 0.22-μm filtered EMM5S media, with coverslips that had been washed in ethanol for 20 minutes and plasma treated for 2 minutes to avoid nonspecific attachment of dyes and other auto-fluorescent particles on the surface. Cells were imaged with an inverted fluorescence microscope (Ti Eclipse, Nikon) equipped with a 60x/1.49 numerical aperture objective (Nikon), illuminated with a 642 nm laser (for imaging SiR647 samples) or 488 nm laser (for imaging mEGFP samples) directed through the objective to achieve TIRF, and recorded with an electron-multiplying charge-coupled device (EMCCD) camera (iXon DU897, Andor). Samples labeled with SiR647 were imaged under low illumination intensity, approximately 20 W/cm^2^. Movies were recorded at a single focal plane near the cell base at 10 frames per second.

### Image analysis and quantification

Image analysis was carried out in the Fiji distribution of ImageJ (Schneider, Rasband, and Eliceiri 2012;Schindelin et al. 2012) and further quantification was performed in Matlab (MathWorks, Inc.), using built-in tools as well as self-written macros and scripts (Supplemental Material). We measured the lengths of filaments in the Average intensity projection of frames 1-5 (AVG1-5) of Pil1p-SiR and Pil1p-mEGFP movies by drawing a line along the full length of visible fluorescence for each filament. We then identified SRAP spots in the Maximum intensity projection of frames 50-200 (MAX50-200), after labeled eisosomes had photobleached. We first generated a preliminary list of SRAP spot positions from the MAX50-200 image by using the Find Maxima command and determining the brightness-weighted centroid of a 3-pixel diameter circle at each point.

For each point in this list, we manually traced the corresponding eisosome filament in the AVG1-5 image with a 3-pixel wide line spanning past the spot position to extend beyond the end of the filament (Figure 2A) and the intensity profile along this line was analyzed in Matlab. Spots which were more than 4 pixels away (280 nm) from the nearest eisosome were discarded (<10% of detected spots). To find the position of the end of the eisosome underlying the diffraction-limited image, the intensity profile was fitted with the following step-like function:

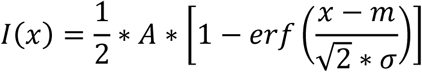

This equation is equivalent to the cumulative intensity of a continuous distribution of Gaussian emitters, where *I(x)* is the intensity along the line coordinate *x, A* is the amplitude, *erf()* is the error function, *m* is the position of the underlying step corresponding to the end of the labeled structure, and *σ* is the standard deviation of the diffraction-limited Gaussian spot. Measured intensity profiles were fitted in Matlab using a nonlinear fitting algorithm, with *A* and *m* as independent variables and *σ* fixed to 1.85 pixels (130nm) representing the diffraction-limited spot width.

We used the PeakFit plugin for FIJI (University of Sussex, http://www.sussex.ac.uk/gdsc/intranet/microscopy/imagej/smlmplugins) to determine super-resolution localizations of the spots that appeared in frames 50 to 200, calibrated with the following parameters: pixel size 70nm, wavelength 642nm, objective NA 1.49, objective proportionality factor 1.4, EM Gain 37.7; resulting in an estimated point-spread function width of 1.837 pixels. This generated a list of localizations with precision <40 nm. From the list of SRAP events' spot centroids determined in the MAX50-200 projection, we matched each SRAP event with all localizations within a 1-pixel radius from the SRAP spot centroid. We calculated the distance from each localization to the fitted eisosome end position projected along the filament line trace (Figure 2A(iv)), and calculated the average distance to the end of all the associated localizations for each SRAP event. For spot centroids that did not have any associated localizations spots of sufficiently high precision, we used the brightness-weighted centroid of the SRAP spot in the projection image to determine its distance from the eisosome end.

To determine the lifetimes of SRAP events, we measured the intensity of a 3-pixel diameter circle centered on the SRAP spot position through the length of the movie and processed these intensity time-traces in Matlab with a Chung-Kennedy filter (Reuel et al. 2012). We computed the lengths of time spans above a threshold intensity, then fit the distribution of lifetimes with a single exponential curve. To estimate the photobleaching rate we measured the mean intensity of an ROI containing an entire cell through the length of the movie. For each ROI's intensity decay profile we subtracted the minimum baseline and normalized the intensities to the maximum value, then computed the average across all movies. We fit the average photobleaching profile with a single exponential curve, starting at frame 5 to avoid biasing the fit with the fast-bleaching autofluorescence component. We estimated the protein unbinding rate by subtracting the bulk photobleaching rate from the SRAP spot lifetime decay rate.

### Characterization of eisosome end localization

We performed simulations to estimate the precision of our method of fitting an error function to the intensity traces of sparsely labeled eisosomes to localize their ends. Indeed, this continuum model might not find the eisosome ends accurately when the structures are only sparsely labeled. Based on published cryo-EM reconstructions of *in vitro* Pil1p filaments, we estimate there are approximately 80 Pil1p proteins per 100nm stretch of eisosome lattice (a hemicylindrical lattice with dimensions roughly as in (Karotki et al. 2011;Kabeche et al. 2011)). Therefore, for the 350-nm (5-pixel) region simulated above we expect 280 possible Pil1p sites, but with our estimated 1 to 3% labeling efficiency there are most likely fewer than 10 fluorescently tagged Pil1p-SiR. We first simulated a number of “emitter positions” in a uniform distribution along a 350-nm line (equivalent to 5 pixels). For 500 simulations for each test case of between 1 and 10 emitters (See Figure 3B and Supplemental Figure S4), we added a Gaussian profile of intensity at each emitter position (mean x_i_, standard deviation 135 nm, peak height of 1 AU) to mimic the point spread function of the microscope, added noise to the traces (random value of mean 0, standard deviation 0.1 AU at each x value), and also added signal from emitters outside the simulated region to account for other fluorophores on the rest of the eisosome body. We fit the resulting intensity profile (10 pixels long, including the 5-pixel region of simulated fluorophores plus 5-pixel tail region) with the error function model described above. We determined the distance from the fitted end position to the true position of the last emitter (position x_0_) and calculated the distance between the position of the last fluorophore (position *x*_o_) and the true end of the eisosome (position 350 nm) in each simulation.

To determine a full population average of these errors, we simulated 5,000 filaments with a distribution of fluorophore numbers based on a binomial distribution (280 trials, 0.01 probability) to determine the probability of any given number of fluorophores on an eisosome. We used a uniform distribution of emitter positions for each filament, generated the intensity profile as above, and fit the error function model to determine the end position. We repeated a similar set of simulations with a single added fluorophore at the end position to account for the artifact of dynamic recovery during the imaging time (Figure 3D and Supplemental Figure S4D).

### Eisosome dynamics model simulations

We compared the distribution of our experimentally measured distances to datasets simulated under different hypotheses. In one model (referred to as the “uniform model”), Pil1p SRAP events occur uniformly along the eisosome, in a second model (referred to as the “end model”) events occur exclusively at the end of the filament, (Figure 2B). For all models, each simulation was initialized by picking one of the eisosome lengths experimentally measured in Pil1p-SiR cells (10,000 runs with each of N = 275 filaments, Supplemental Figure S1C). For the uniform model, the true SRAP spot positions were simulated by picking a number following a uniform distribution between zero and half the filament length; for the end model, the true SRAP spot position was taken as the true end position of the eisosome end (position 0) and a number following a Gaussian distribution (mean 0, standard deviation 25 nm) was added to represent the spot localization with uncertainty as measured experimentally (Figure 2B-C). For each simulation, we added a number following a Gaussian distribution with mean 0 and standard deviation 120 nm to the true position of the eisosome end (position 0) to simulate the localization precision of the experimental fit of the eisosome end in our image analysis (as described above and in Supplemental Figure S4E). The position of the eisosome ends and SRAP spots were then subtracted from each other to determine the relative positions of the SRAP spots. In a second set of simulations to account for the fitting bias arising from a dynamic filament end, we used for the eisosome end position distribution a Gaussian distribution with mean −90 nm and standard deviation 94 nm (as in Figure 2D).

We also simulated a third class of models (referred to as “hybrid models”) where events occur uniformly within a zone of defined length (e.g. 200 nm) at the eisosome end. For the hybrid models, the true SRAP spot position was simulated by picking a number following a uniform distribution between zero and the length of the end zone (e.g. 200 nm), and the end position and noise terms were generated with unbiased Gaussian distributions as described above.

## Acknowledgements

We thank Dr. Ronan Fernandez for assistance in creating yeast strains, and members of the Berro lab for helpful discussions. This research was supported in part by the NIH/NIGMS grant R01GM115636. MML was supported by the NIH training grant T32GM008283. We also acknowledge support from the Raymond and Beverly Sackler Institute for Biological, Physical and Engineering Sciences at Yale University.

## Supplementary Information

**Supplemental Figure S1.**
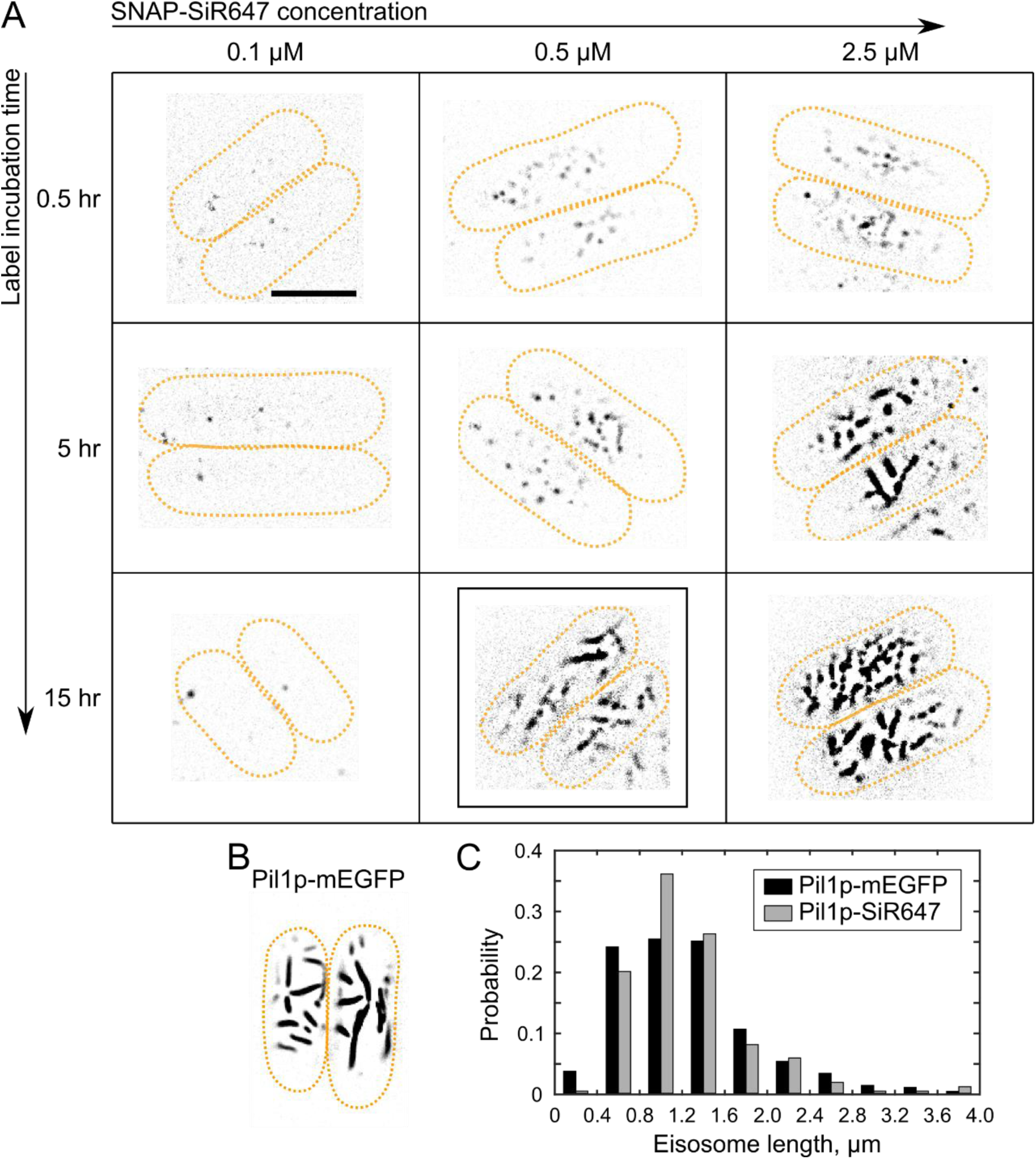
SNAP labeling of Pil1p in live fission yeast cells. (A) *S. pombe* cells expressing Pil1p-SNAP were incubated with SNAP-SiR647 at indicated concentrations in EMM5S media for various times, washed and imaged in TIRF. The boxed panel highlights the sample condition used for further imaging and analysis, 15 hours at 0.5 μM SNAP-SiR647. (B) Cells expressing Pil1p-mEGFP imaged in TIRF. All image panels are at the same length scale with scale bar 5 μm, and same brightness scale (except for B). (C) Histograms of eisosome lengths measured in cells with Pil1p-SiR (grey, 1240 ± 580 nm for N = 275 eisosomes measured in 20 cells) or Pil1p-mEGFP (black, 1250 ± 650 nm for N = 304 eisosomes measured in 22 cells) show no significant difference by two-sample Kolmogorov-Smirnov test (*p* = 0.33).

**Supplemental Figure S2.**
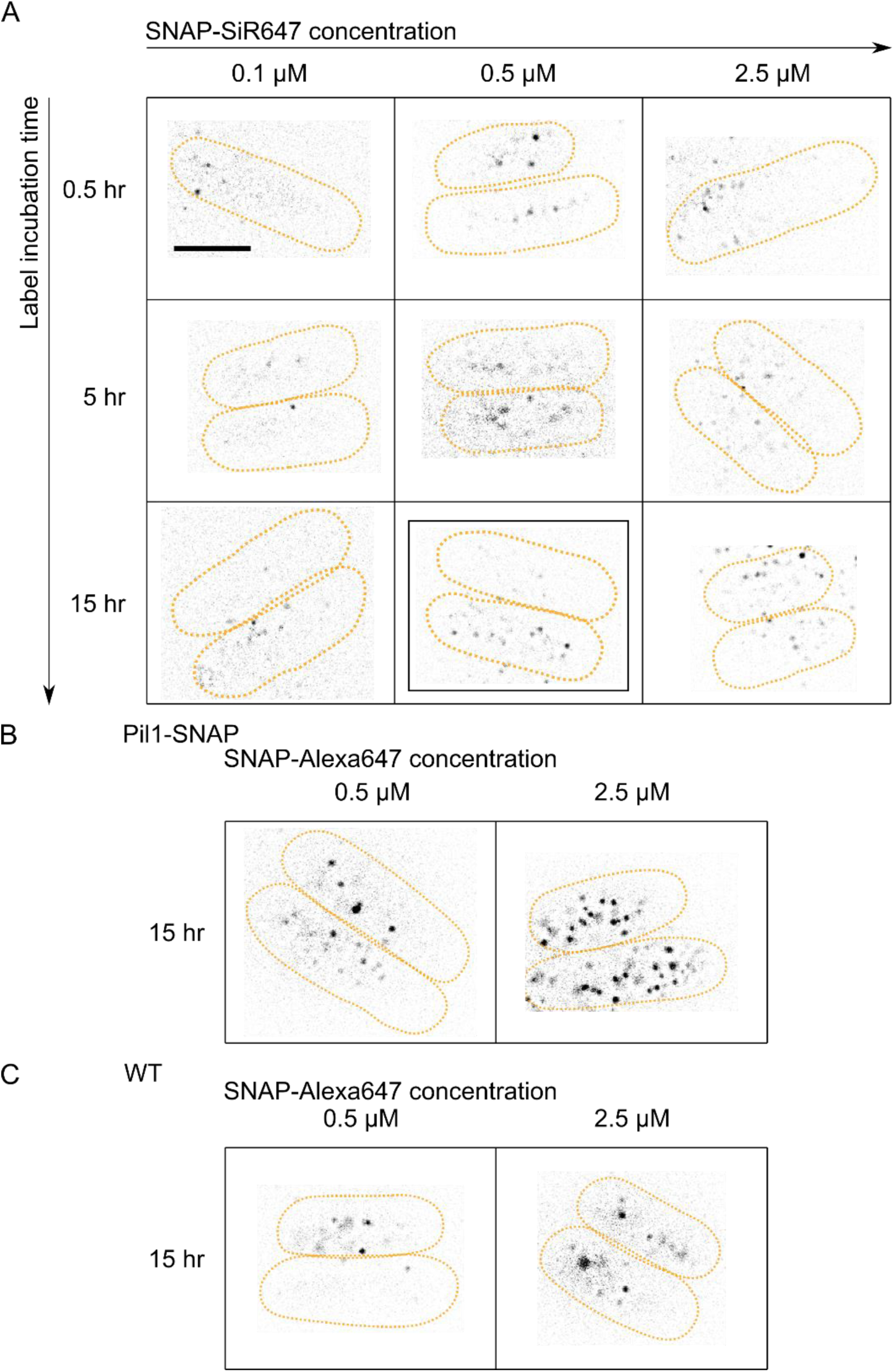
Comparison of SNAP labeling and nonspecific binding. (A) Wild-type cells were incubated with SNAP-SiR647, washed, and imaged in TIRF as described in the text. (B and C) Cells expressing Pil1p-SNAP (B) or wild-type cells (C) were incubated with SNAP-Alexa647, washed, and imaged as described. Images shown are inverted contrast, Maximum intensity projections of 20-sec movies with median-filter background subtracted. Cell outlines are drawn in orange dash; all image panels are at same scale with scale bar 5 μm.

**Supplemental Figure S3.**
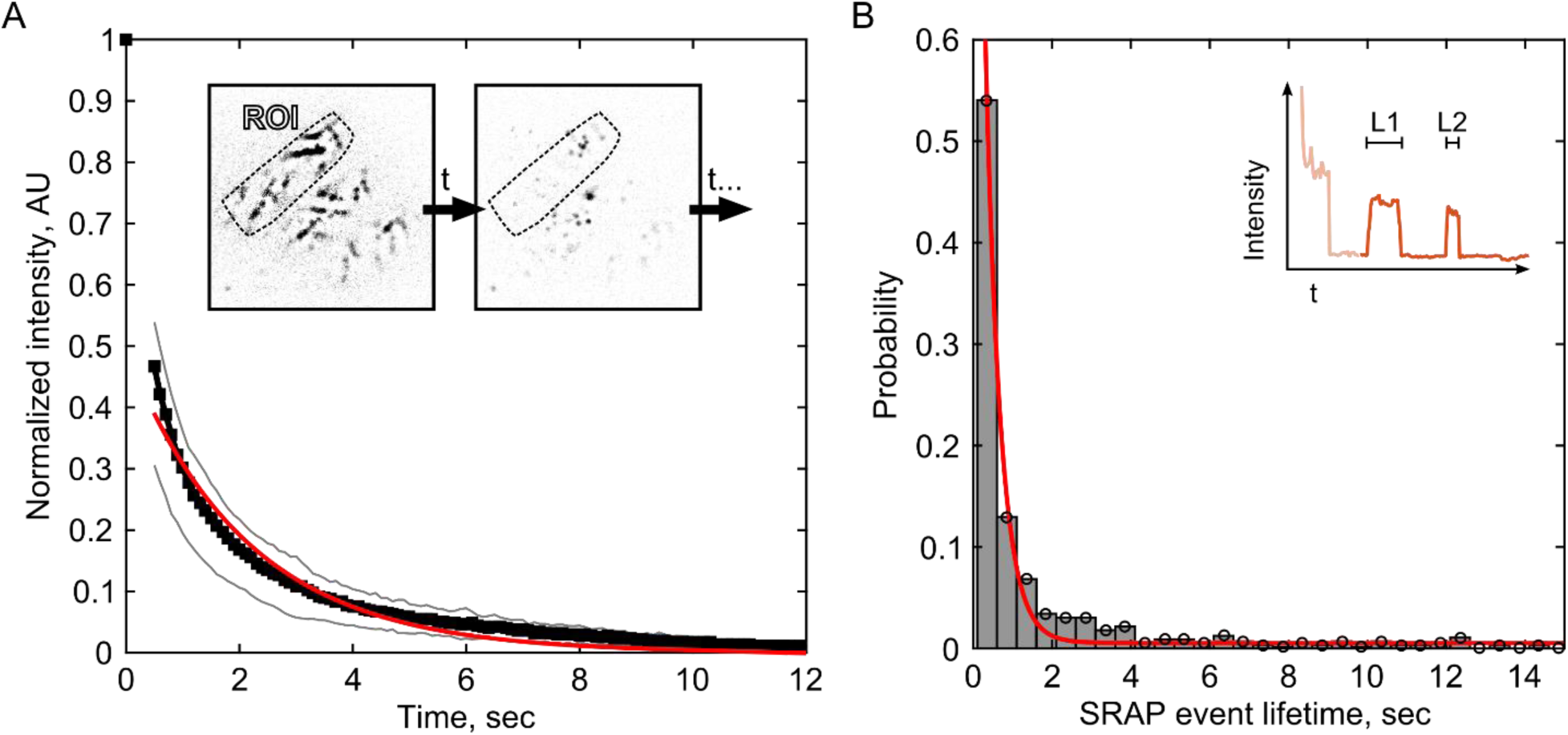
Comparison of photobleaching rate and SRAP event lifetimes. (A) Photobleaching was measured as fluorescence intensity of ROIs drawn around cells throughout the movie stack. For each curve the baseline was subtracted and curves were normalized to their initial value, then all ten curves were averaged (black squares, gray lines +/− standard deviation). The average curve (starting at frame 5) was fit with a single exponential (red curve) to determine the decay rate of −0.48±0.03 sec^−1^. Inset: schematic of ROI measured through movie stack. (B) Distribution of lifetimes of fluorescence events at SRAP spots. The distribution (N=558 events, 433 longer than 2 frames) was fitted with a single exponential (red curve) to determine the off-rate of −2.62±0.22 sec^−1^. Inset: schematic showing an example fluorescence intensity trace with multiple events.

**Supplemental Figure S4.**
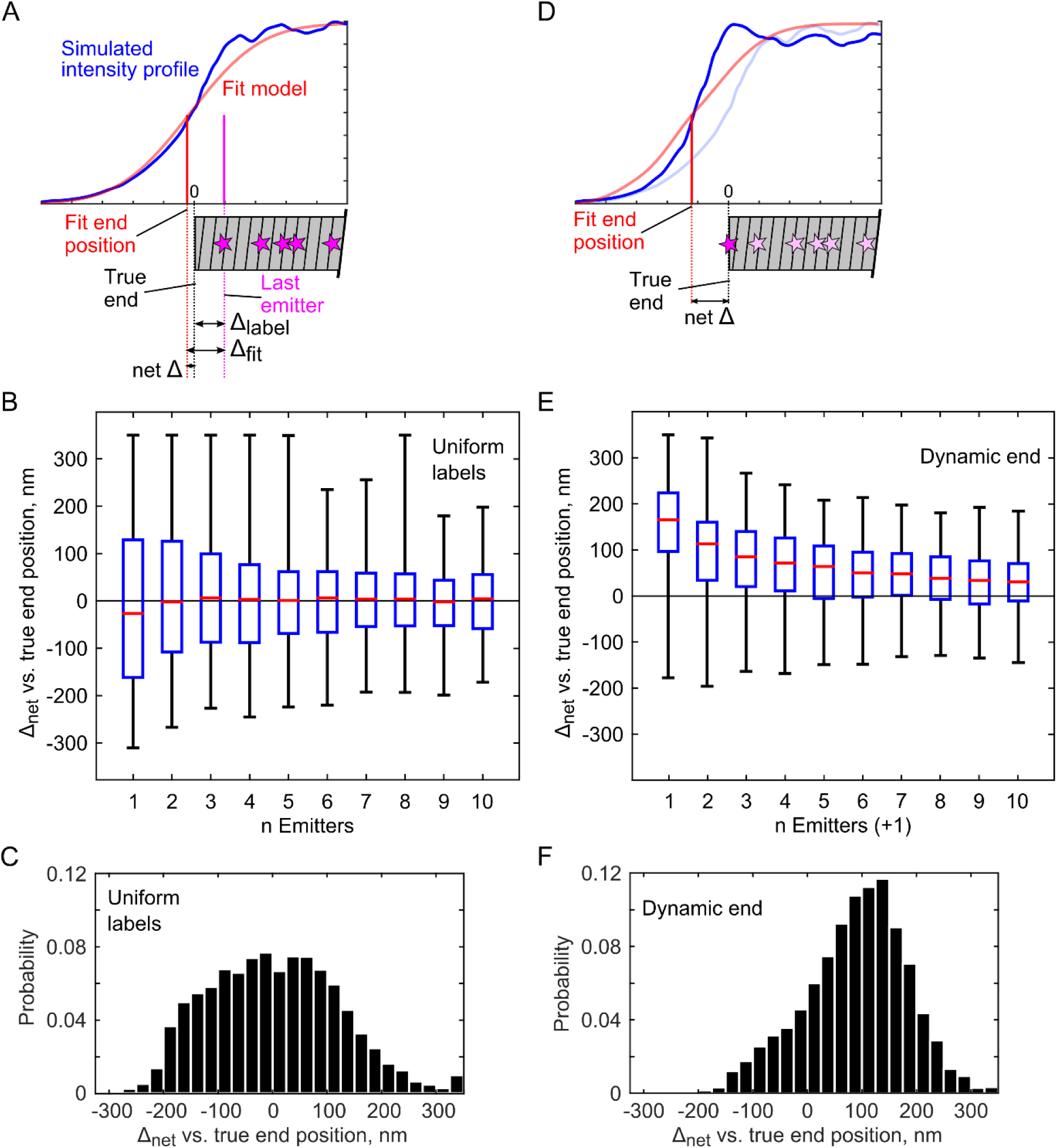
Characterization of errors in simulated eisosome end localizations due to sparse labeling. (A) Schematic of errors in fitting: mock fluorescence intensity traces (blue) were generated by simulating discrete numbers of emitters in a 350-nm region (5 pixels) and then fit with the error function model (red). Legend: Δ_label_, distance between the last emitter (closest to the eisosome end) and the eisosome end; Δ_fit_, distance between the eisosome end position estimated by fitting the fluorescence intensity and the last emitter; Δ_net_, difference between the eisosome end position estimated by fitting the fluorescence intensity and the true end position. (B) Difference between the fitted end position and the true end of the simulated eisosome (Δ_net_), calculated for discrete numbers of emitters (N=500 simulations for each n Emitters). (C) Distribution of the differences between the fitted end and the true end (Δ_net_) for a population of eisosome traces simulated according to 1% labeling. Δ_fit_ and Δ_label_ nearly balance each other, with Δ_net_ average 3.2 ± 119 nm S.D. (N = 5,000 simulations). (D) To determine the effect of recruiting new fluorescent molecules only at the filament ends, mock fluorescence intensity traces (blue) were generated by simulating a discrete number of emitters in a 350-nm region with one additional emitter at the eisosome end, and then fit with the error function model (red). The intensity profile is skewed beyond the true eisosome end position and Δ_label_ is reduced to zero. (E) Difference between the fitted end position and the true end of the simulated eisosome filament (Δ_net_), calculated for discrete numbers of emitters with an additional emitter at the eisosome end (N = 500 simulations for each n Emitters). (F) Distribution of the differences between the fitted end and the true end (Δ_net_) for a population of eisosome traces simulated according to 1% labeling with a single extra emitter added to the eisosome end, Δ_net_ average 89.3 ± 94 nm S.D. (N=5,000 simulations). Box plots show median (red line), 25th and 75th percentile (blue box), and farthest outliers (whiskers). In all plots, the sign of Δ is given as the effect on the calculated spot position (i.e. estimating the end to be past the structure causes the calculated SRAP spot position to be shifted toward the filament interior, a positive value).

**Supplemental Figure S5.**
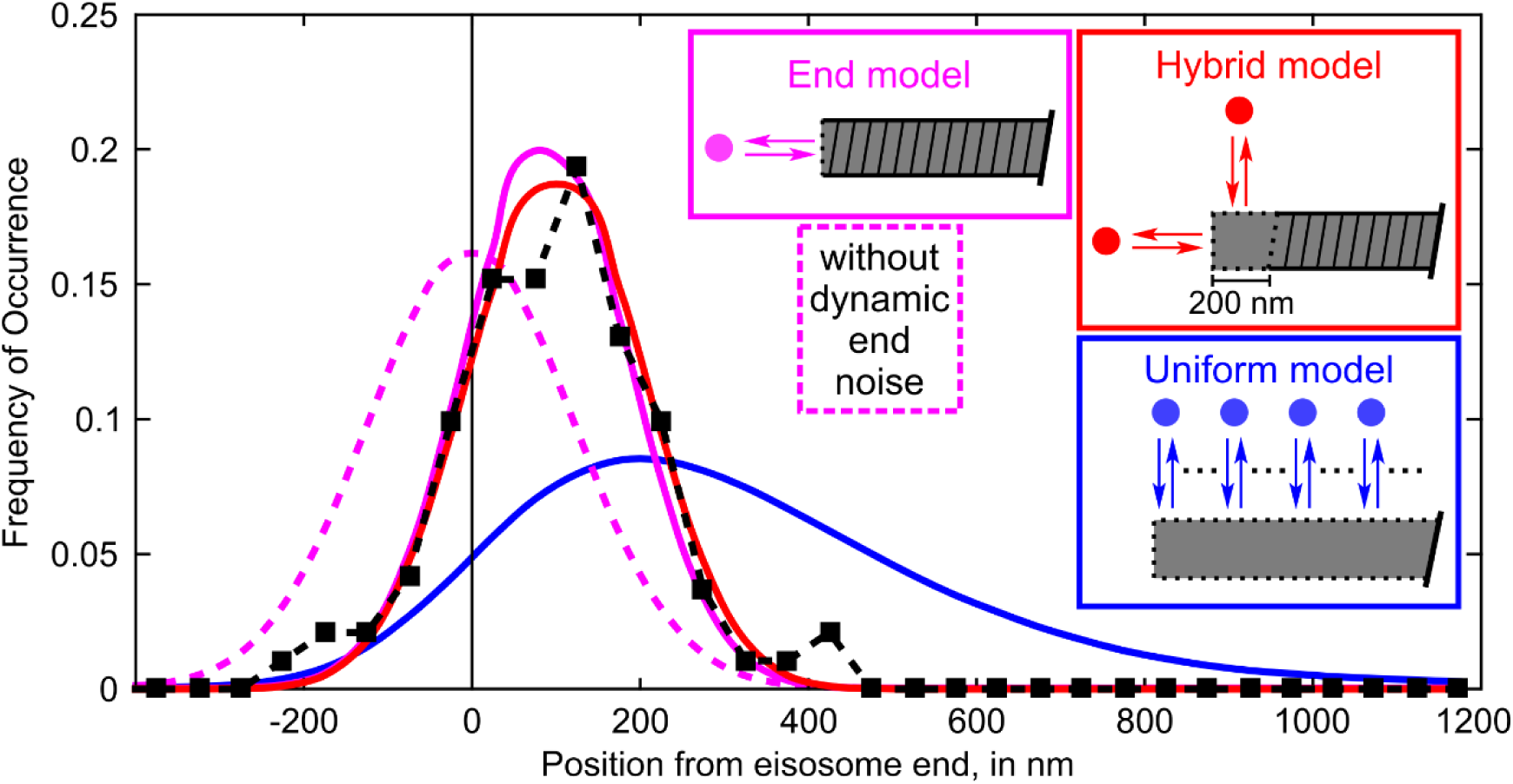
Alternate model for eisosome recovery dynamics. Probability distributions of measured distances (black squares, N = 191 spot/filament pairs across 20 cells) and simulation results (similar to Figure 3, N = 275,000 runs for each tested model). An additional “hybrid” model is shown (red), where simulations used simple Gaussian noise for the eisosome end position and calculated spot positions uniformly distributed within a 200-nm zone at the eisosome end. Simulated end model with biased (solid magenta) and unbiased (dashed magenta) end noise, uniform model (blue), and measured dataset (black) are as shown in Figure 3. The hybrid model average position is 100 ± 103 nm. The difference between measured data (97 ± 119 nm) and the simulated hybrid model is not statistically significant, by two-sample Kolmogorov-Smirnov test (*p* = 0.32)

**Table S1.**
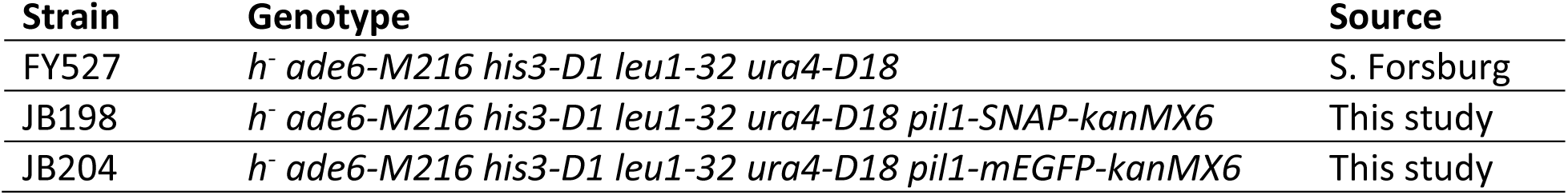
*S. pombe* strains used in this study

| Strain | Genotype | Source |
| --- | --- | --- |
| FY527 | *h<sup>-</sup> ade6-M216 his3-D1 leu1-32 ura4-D18* | S. Forsburg |
| JB198 | *h<sup>-</sup> ade6-M216 his3-D1 leu1-32 ura4-D18 pil1-SNAP-kanMX6* | This study |
| JB204 | *h<sup>-</sup> ade6-M216 his3-D1 leu1-32 ura4-D18 pil1-mEGFP-kanMX6* | This study |

**Supplemental Movie S1**

Cells expressing Pil1p-SNAP were labeled with SNAP-SiR647 at 0.5 μM for 15 hours, washed and imaged in TIRF. Movie was recorded at a single focal plane near the cell base at 10 frames per second. This movie file was used to generate parts of Figure 1 and Supplemental Figure S1. Scale bar 5 μm

**Scripts for image analysis and quantification**

**File 1: ImageJ Macro for finding, measuring recovery events and tracing eisosome ends:** Supplement_script_ij.txt

**File 2: Matlab script for fitting of eisosome filament ends, calculation of distance from SRAP spots to eisosome ends, simulations of eisosome end intensity profiles, and simulations of eisosome dynamics models:** Supplement_script_matlab.txt

The results tables saved from ImageJ macro analysis can be opened in Excel to edit columns and clean to remove headings. The results tables for each movie file are copied into a separate Matlab file that loads the results tables as multidimensional arrays, called `f []` and `c []`, and the results of the GDSC SMLM PeakFit plugin, called `smlm []`.

A separate results file is needed containing line measurements of eisosome lengths, manually drawn in AVG1-5 projections from all movies, `total []`.

